# Structural Basis for Autoinhibition by the Dephosphorylated Regulatory Domain of Ycf1

**DOI:** 10.1101/2023.06.22.546176

**Authors:** Nitesh Kumar Khandelwal, Thomas M. Tomasiak

## Abstract

Yeast Cadmium Factor 1 (Ycf1) sequesters glutathione and glutathione-heavy metal conjugates into yeast vacuoles as a cellular detoxification mechanism. Ycf1 belongs to the C subfamily of ATP Binding Cassette (ABC) transporters characterized by long flexible linkers, notably the regulatory domain (R-domain). R-domain phosphorylation is necessary for activity, whereas dephosphorylation induces autoinhibition through an undefined mechanism. Because of its transient and dynamic nature, no structure of the dephosphorylated Ycf1 exists, limiting understanding of this R-domain regulation. Here, we capture the dephosphorylated Ycf1 using cryo-EM and show that the unphosphorylated R-domain indeed forms an ordered structure with an unexpected helix-strand hairpin topology bound within the Ycf1 substrate cavity. This architecture and binding mode resemble that of a viral peptide inhibitor of an ABC transporter and the secreted bacterial WXG peptide toxins. We further reveal the subset of phosphorylation sites within the hairpin turn that drive the reorganization of the R-domain conformation, suggesting a mechanism for Ycf1 activation by phosphorylation-dependent release of R-domain mediated autoinhibition.

## Introduction

Yeast Cadmium Factor 1 (Ycf1) belongs to the ATP Binding Cassette (ABC) transporter family and sequesters glutathione and glutathione heavy metal conjugates to the vacuole to detoxify reactive oxygen species and heavy metals (Cd^2+^, Se^2+^, Hg^2+^, As^2+^)^1–4^. In eukaryotes, ABC transporters are divided into five subfamilies (A, B, C, D, and G) with different topologies built around a core architecture made of two nucleotide-binding domains (NBDs) where ATP is hydrolyzed and two transmembrane domains (TMDs) through which substrate is transported. Ycf1^5,6^ belongs to the C subfamily (termed ABCC),^7,8^ which, in addition to the core architecture, includes several evolutionary additions, including an additional TMD (called TMD0), a lasso domain that links TMD0 and TMD1, and a flexible linker domain called the regulatory domain (R-domain) that connects NBD1 and TMD2^9^.

R-domains play prominent roles in transporter activation and have unique properties, making them hubs of signaling integration^10^. They are typically large (∼60-200 amino acids), and their phosphorylation is indispensable for activity in Ycf1 and the cystic fibrosis trans-membrane regulator (CFTR^6,11,12^, a related chloride transporter that causes cystic fibrosis when mutated^13^ and can also transport glutathione^14^. R-domains connect NBD1 to TMD2, placing them near several key allosteric and catalytic elements, such as the ATP binding sites. In Ycf1, mutation of three R-domain sites (S903, S908, and T911) results in loss of survival in the presence of cadmium, a Ycf1 substrate^12^. CFTR contains a more extensive R-domain with six activity-driving phosphosites and 20 disease-associated mutation sites. In CFTR, activity also depends on R-domain phosphorylation driven by interactions with Protein Kinase A and other kinases such as PKC. It is also the site of several protein-protein interactions^10^. In Multidrug Resistance Protein 1 (MRP1), another glutathione and glutathione conjugate ABCC transporter^15^ and proposed ortholog of Ycf1^16^, the R-domain also controls relative substrate selectivity for the heavy metals selenium or arsenic^17^.

Conflicting reports detail how the R-domain structure transitions between phosphorylated and dephosphorylated states. It has been proposed that the R-domain is disordered mainly because of low sequence complexity dominated by repeats of negatively charged amino acids thought to have derived from an intronic evolutionary origin in CFTR^18^. This conclusion is supported by several biophysical investigations in isolated R-domains by NMR^19^, circular dichroism^20^, protease accessibility^20^, and SDS-PAGE migration assays that show little secondary structure and suggest substantial intrinsic disorder. They also imply that the R-domain mainly resides between NBD1 and NBD2 when unphosphorylated, thereby poised to play an autoinhibitory role^21^. In this model, the R-domain mobility appears to become even more diffuse upon phosphorylation and is subsequently released from its position between the NBDs to relieve autoinhibition and permit activity.

In contrast to these findings from isolated segments, investigations in full-length transporters, including cryo-EM, provide evidence that more well-ordered intermediates of the R-domain might exist. For example, although the R-domain is completely unresolved in cryo-EM structures of dephosphorylated CFTR, they do show diffuse electron density for the R-domain oriented between both halves of CFTR,^7^. On the other hand, phosphorylated CFTR structures show limited R-domain segments built only as c-alpha traces because of poor resolvability, nevertheless suggesting interactions along the periphery of NBD1. More recently, high-resolution Ycf1 cryo-EM structures clarify this region and reveal the topology of a well-resolved R-domain and its residues bound to NBD1^5,6^. In addition, our phosphorylated Ycf1 structure directly shows cryo-EM density for a trio of phosphorylated residues (S908, T911, and S914) that directly engage basic surfaces within this region^6^. Finally, surface accessibility studies on full-length CFTR^22^ and Ycf^5^ suggest that the dephosphorylated R-domain engages a significant portion of the TMDs, while the phosphorylated R-domain only engages NBD1. Altogether, these structures point to an R-domain architecture far more structured than previously alluded to by biophysical investigations.

Here, we report the cryo-EM structure of Ycf1 in a dephosphorylated state that captures key allosteric and orthosteric interactions. Changes in substrate-induced ATPase acceleration, cell survival on toxic heavy metal cadmium, and changes in accessibility to proteases support a new model for the dephosphorylated R-domain inhibitory mechanism as well as insights into ABCC family transporter activation.

## Results

### Dephosphorylated Ycf1 resembles phosphorylated IFwide and lacks the R-domain around NBD1

Many ABCC family transporter family members are regulated by dynamic phosphorylation of an intrinsically disordered R-domain^9^. Our previous cryo-EM structure of Ycf1 revealed a partially structured conformation of the phosphorylated R-domain engaged with NBD1 that, upon dephosphorylation with alkaline phosphatase, drastically lowered basal ATPase activity and caused changes to the Ycf1 architecture^6^. Here, we sought to determine the structural changes in dephosphorylated Ycf1-E1435Q expressed and purified from the *Saccharomyces cerevisiae* DSY5 strain. We generated the dephosphorylated state by treating purified phosphorylated Ycf1 with alkaline phosphatase, followed by A subsequent round of size exclusion purification to yield a highly homogenous dephosphorylated sample (Figure 1A - top) verified by a phosphoprotein dye (Figure 1A – bottom).

**Figure 1.**
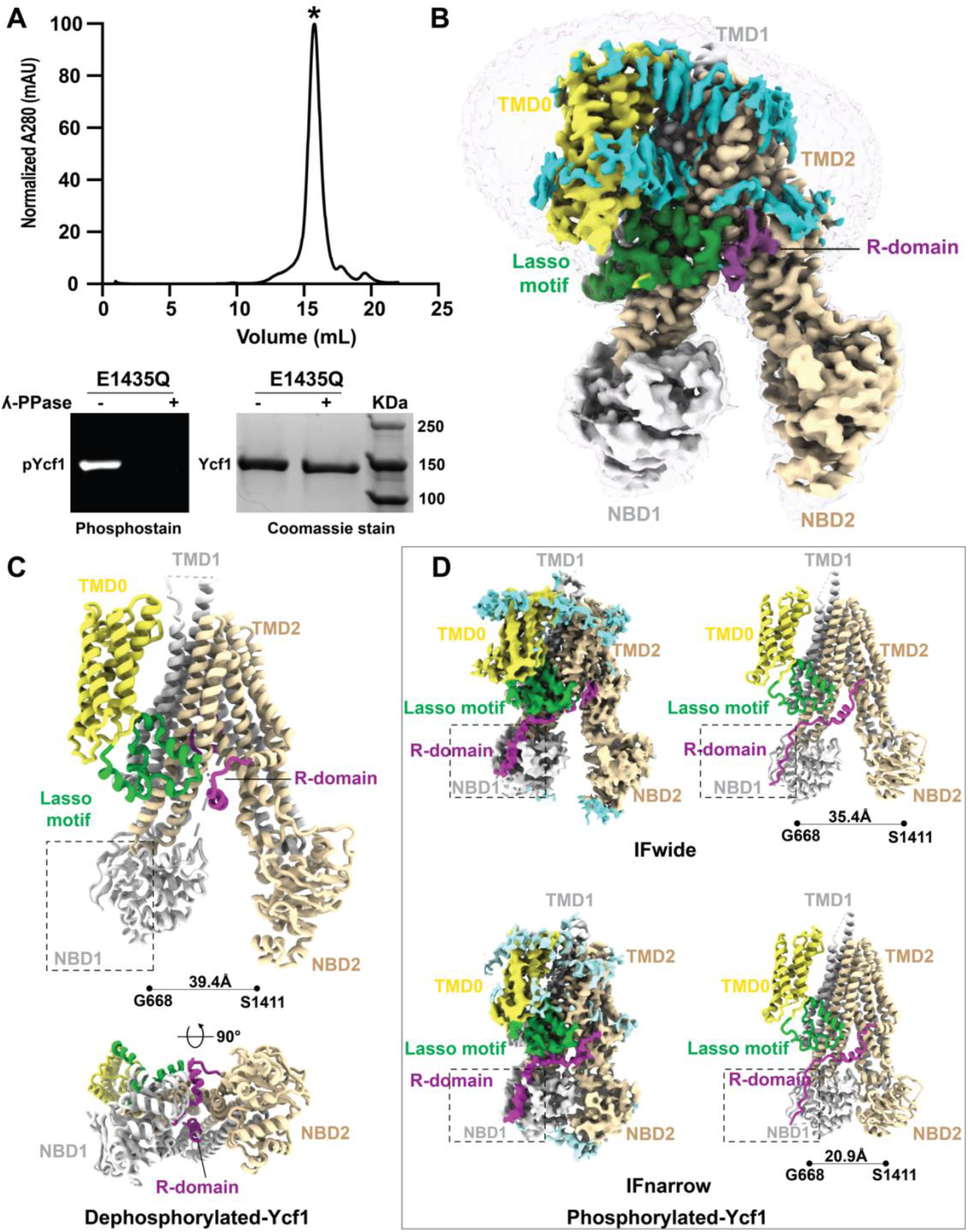
Structure determination of dephosphorylated Ycf1. **A**. Size exclusion chromatography profile of dephosphorylated Ycf1. **B**. Cryo-EM density of the dephosphorylated Ycf1 structure. TMD0 is colored yellow, TMD1 and NBD1 are colored white, TMD2 and NBD2 are colored wheat, the lasso domains are colored green, the regulatory domain (R-domain) is purple, and ordered lipids are colored light blue. **C** Cartoon of the dephosphorylated Ycf1 structure. **D**. Cryo-EM density and model of phosphorylated Ycf1 structures IFwide (PDBID: 7M69) and IFnarrow (PDBID:7M68) colored as in B. The area of missing density for the R-domain around NBD is highlighted with a hashed box.

The homogenous sample enabled cryo-EM structure determination of dephosphorylated Ycf-E1435Q to an overall final resolution of 3.1 Å Figures 1B-C and Figure S1), a higher resolution than previously published phosphorylated structures (3.1Å vs. 3.2Å,3.4Å, and 4.0Å). Surprisingly, dephosphorylation of Ycf1 leads to a complete loss of the R-domain density at its docking sites on NBD1 (Figure 1B-C) previously observed in the phosphorylated structures (Figure 1D). Instead, considerable new electron density was visible within the substrate-binding cavity between TMD1 and TMD2. This density displayed several continuous segments that allowed for the structural assignment of the C-terminal half of the R-domain (residues 895-935) continuous with TMD2 (Figure 2A, see Methods for details of structure assignment).

**Figure 2.**
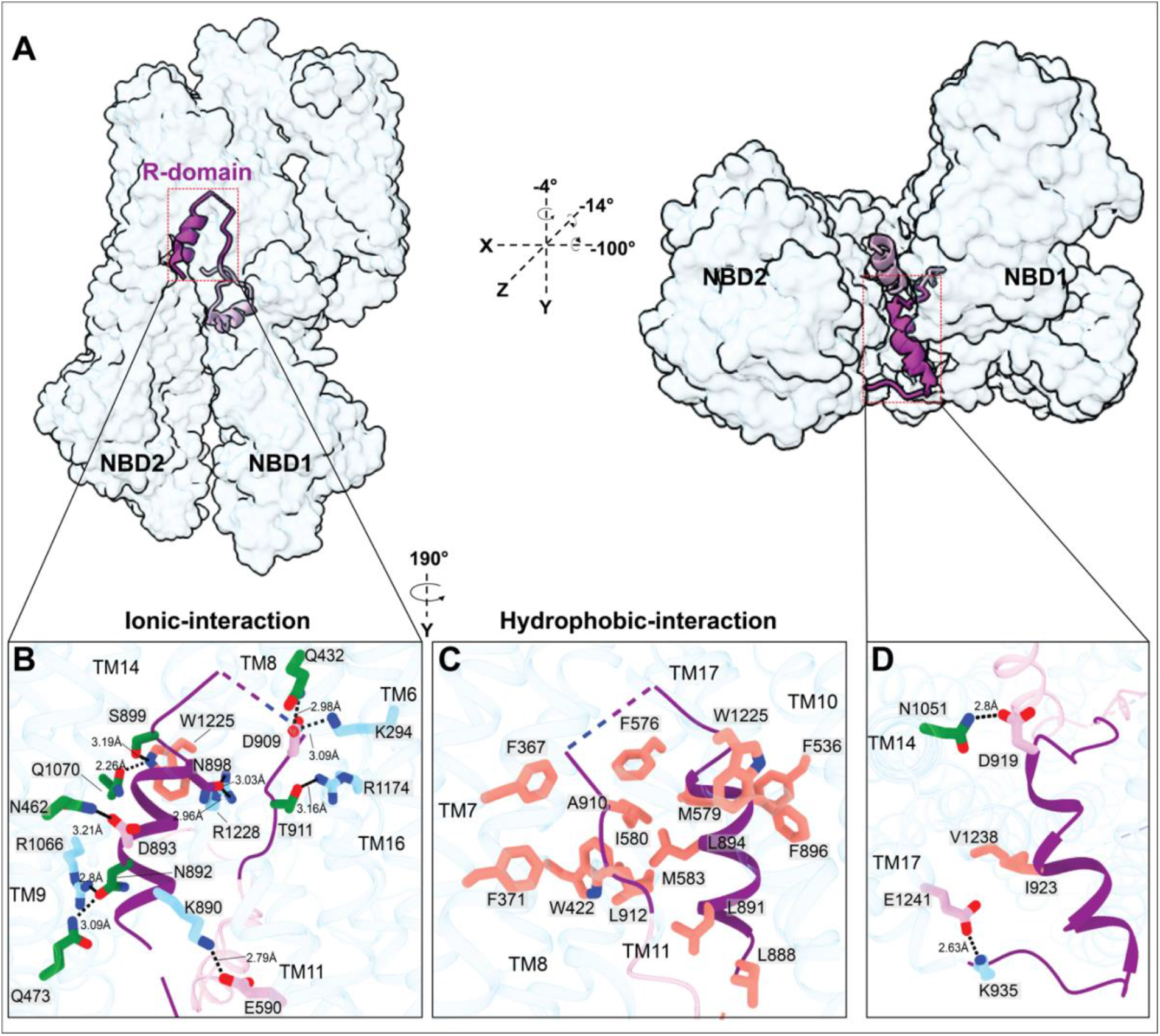
Representation of the R-domain in the Ycf1 substrate cavity. **A**. A space-filling model of Ycf1 with a transverse section of the TMD removed to show the R-domain from a side view (left) and bottom view of the NBDs (right). **B**. Electrostatic interactions between the R-domain and the substrate binding cavity. The R-domain backbone is colored purple, with interacting residues and the Ycf1 backbone colored white. Amino acid side chains interacting between the R-domain and Ycf1 are colored based on physical amino acid properties (Hydrophobic residues are colored orange, positively charged amino residues are in cyan, negatively charged residues are in pink, and polar-uncharged residues colored in green. **C**. Hydrophobic interactions with the same color scheme as in B. Detailed view of R-domain hydrophobic interactions with the TMD region of Ycf1 and. **D**. the C-terminal region of the R-domain.

Apart from the substantive gross R-domain rearrangement, the dephosphorylated Ycf1 architecture remains almost unchanged from the IFwide conformation of phosphorylated Ycf1 and exists in a single major conformation (compared to three conformations observed in our dataset of the phosphorylated state (Figures 1B-D)^6^. We could successfully build all 17 transmembrane helices (TMHs), the lasso region, both NBDs and much of the R-domain in its new conformation. In addition, several phospholipids are peripherally bound to the transmembrane region embedded in the detergent micelle (Figure 1B-C and Figure S2). Most of the Ycf1 structure was of higher quality (as judged by map quality and correlation coefficient) than previous phosphorylated structures, as reflected by the higher resolution (Figure S1B).

### Dephosphorylation leads to R-domain occlusion of the substrate cavity

In our structure, the R-domain directly interacts with the substrate binding cavity (Figure 1C bottom). It adopts an unexpected helix-strand hairpin motif constructed of an alpha-helix (residues 888-900) with pronounced corresponding density in the cryo-EM map. Following this helix, the R-domain is capped by a loop packed against a strand (residues 909-914) that eventually connects to a helix leading to TMD2 (Figure 2A-C). Gaps in the cryo-EM maps at the turn of the hairpin (residues 900-908) precluded structure assignment in these regions, which coincides with two phosphorylation sites (S903 and S908) (Figure 2A-C). Nevertheless, the arrangement of the visible secondary structure approximately matches those from Alphafold predictions of the R-domain (Figure S3), which proposes an alpha-helical conformation of residues 883-896 engaged with the distal surface of NBD1. The presence of this helix in the phosphorylated Ycf1 structure was noted but could not be modeled in that structure^6^.

The tightly packed R-domain extensively buries the hydrophobic surface of the TMDs, (Figure 2B-C), ∼1800Å of solvent-exposed surface area, and forms several electrostatic interactions. Within the R-domain, the hairpin buries several pairs of hydrophobic residues within itself, including Leu894 against Leu912 and Leu894 against Ala910, analogous to a leucine zipper motif (Figure 2C and Figure S3B). These interactions also make hydrophobic contacts with residues of the substrate cavity (Phe576 - Trp422, Met579, Ile580, Met 583, and Trp1225). The R-domain is further stabilized by polar and H-bonding contacts, including Asn892 to Gln473 and Arg1066, Lys890 to Glu590, Asp893 to Asn462, Asn898 to Arg1228, and Ser899 to the indole nitrogen of Trp422 (Figure 2B). In addition, critical electrostatic contacts form between Asp909 and a bundle of positive residues that includes Lys294 and Gln432. Finally, Thr914 forms a pair of interactions; a polar interaction between its g-hydroxyl and Arg1174 and a second hydrophobic interaction between its g-methyl and Phe367. At the C-terminal region of the R-domain, a second helix formed by residues 920-935 and capped by two glycines moves as a rigid body to form several electrostatic interactions (Figure 2D). Asp919 and Lys935 form H-bonds with Asn1051 and Glu1241, respectively, while Ile923 and Val1238 make nonpolar interactions. Finally, the interactions in the substrate cavity accompany dephosphorylated R-domain contacts with a critical allosteric junction, the “GRD” motif of ICL-1, which couples with the X-loop of NBD1 to communicate ATP binding^23^.

### The R-domain contacts resemble MRP1-substrate and TAP1/TAP2-inhibitor interactions

The new position of the dephosphorylated R-domain occludes the substrate binding site through an extensive interaction network, reducing the cavity volume from 7123Å^3^ in the phosphorylated state to 3934Å^3^ (Figure 3A-B). Although a substrate-bound state of Ycf1 is unavailable, this positioning of the R-domain mimics the binding mode of the glutathione moiety of the substrate Leukotriene C4 (LTC4) in the structure of bovine MRP1^8^, a close functional homolog of Ycf1^16^. The glutathione moiety interacting residues in MRP1 are conserved in Ycf1 and adopt the same spatial positions in the Ycf1 structure (Figure 3C). Several residues of the R-domain and substrate cavity contacts are also conserved in the two structures (Figure 3D-E), including electrostatic interactions with Lys294, His297, R1174, and R1228 (Figure 3D). In addition, several hydrophobic sites, such as the Phe367, Trp422, and Phe576, form hydrophobic interactions with several conserved R-domain elements.

**Figure 3.**
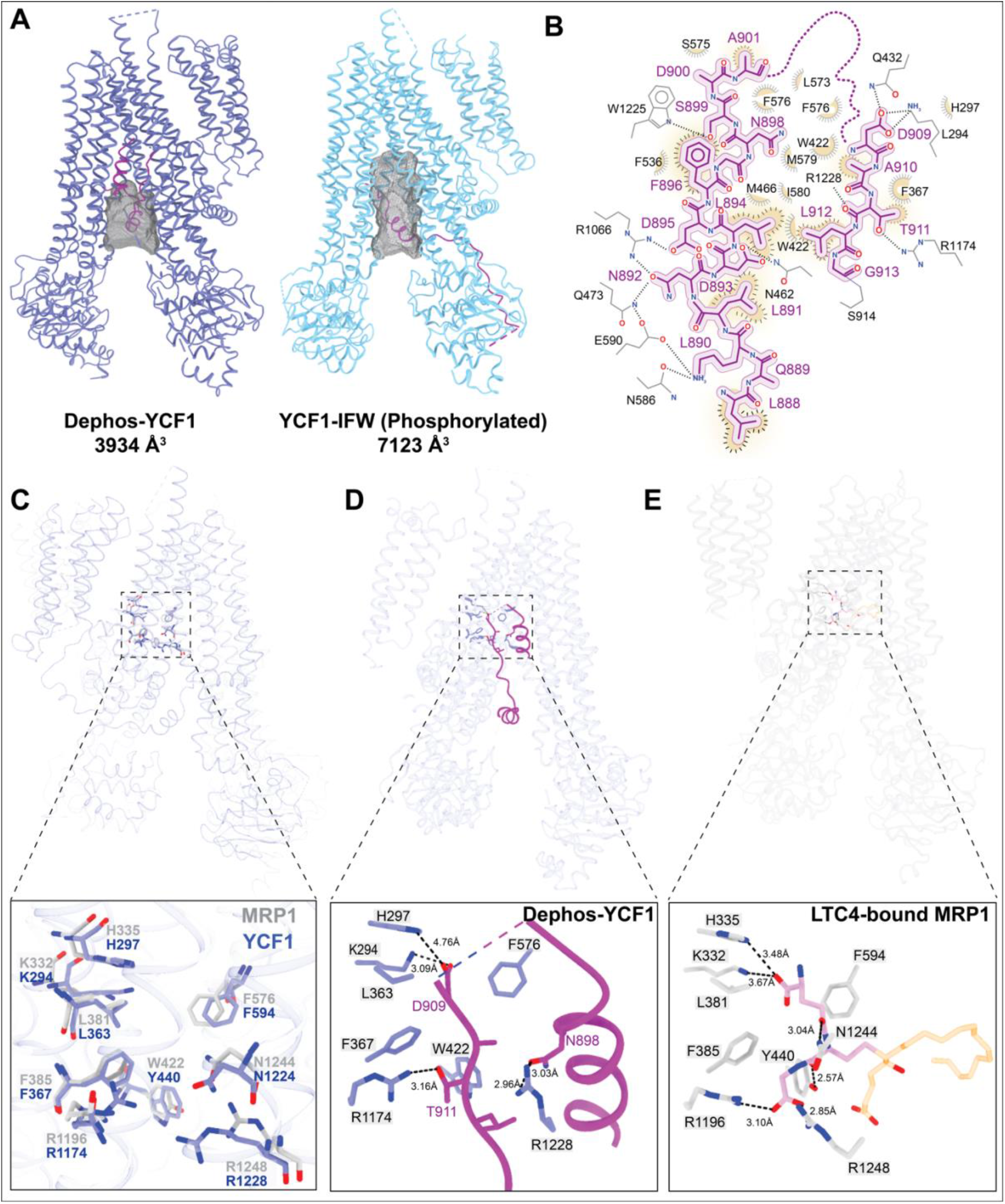
Comparison of transport substrate binding pocket in dephosphorylated Ycf1, phosphorylated Ycf1, and substrate-bound MRP1. **A**. Solvent accessible space-filling model of the Ycf1 substrate cavity. **B**. Ligplot detailing specific interactions between dephosphorylated Ycf1 and the R-domain in its binding cavity. **C**. Comparisons of the phosphorylated Ycf1 R-domain binding sit to MRP1 (PDB IB: 5UJA^8^). **D**. Detailed view of the dephosphorylated Ycf1 R-domain interaction in the substrate binding pocket. **E**. Interaction network of the glutathione moiety in Leukotriene C4 bound to MRP1 for comparison (PDB ID: 5U1D^27^).

Strikingly, alignment of the R-domain structure with the previous structure of the peptide presenting antigen transporter TAP1/TAP2 bound to an inhibitor from herpes simplex virus^27^ reveals unexpected similarities and structural overlap (Figure S4). The ICP47 peptide exhibits an identical architecture with nearly equivalent contacts throughout the substrate cavity. Surprisingly similar elements are visible in the R-domain and the ICP47 peptide, including a similar helical hairpin near hydrophobic residues. These are formed by Trp3, Met7, Phe11, Tyr22, Val25, and Ile29 that are stabilized by a large hydrophobic pocket in TAP1/TAP2. At the hairpin connection in TAP1/2, a sizeable basic pocket resembling Ycf1 forms and is comprised of Arg381(B), Arg380(B), and His 384(B).

### R-domain hairpin phosphorylation of S908 and S903 mainly drives activity

The structural results observed are conducive to a state that blocks enzymatic activity. Basal ATPase activity is strongly inhibited by dephosphorylated Ycf1 and by R-domain mutants, as previously shown^6^. Furthermore, GSSG-dependent stimulation of ATPase activity observed in WT Ycf1 is completely lost in both dephosphorylated Ycf1 and the S908A R-domain mutant. Interestingly, the S903A mutation does affect GSSG-potentiated ATPase activity at 8 µM (EC_50_) concentration, whereas partial induction is observed in the T911A variant (Figure 4A). Notably, phosphosites S903 and S908, which severely impact ATPase activity when mutated, could not be resolved in the dephosphorylated cryo-EM map. These results suggest that either 1) glutathione binding is blocked or 2) glutathione binding is not allosterically communicated with the ATPase sites in these mutants.

**Figure 4.**
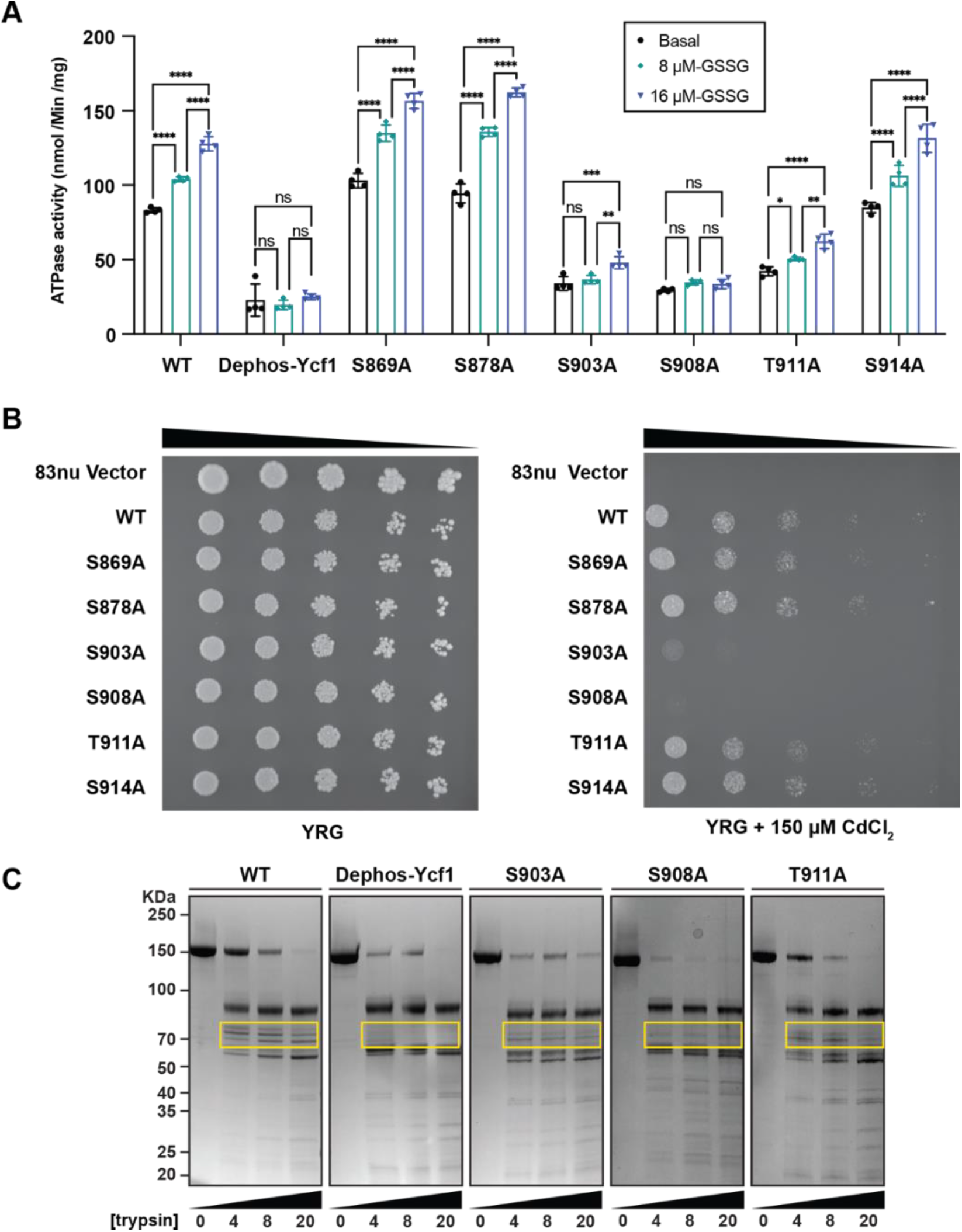
Functional characterization of Ycf1 activity upon dephosphorylation and mutation. **A**. ATPase rates in different Ycf1 phosphorylation states and Ycf1 mutants. ATPase rates of Ycf1 with and without glutathione acceleration. All values are the mean of four replicates ± standard deviation (S.D.) **B**. Spot assays showing survival of cells with Ycf1 mutants. **C**. Limited proteolysis of phosphorylated Ycf1, dephosphorylated Ycf1, and R-domain mutants.

To measure the overall physiological effect of dephosphorylation on substrate translocation, we performed a survival assay in which Ycf1-dependent transport is necessary for yeast growth on media containing 150 µM CdCl_2_, a toxic heavy metal substrate of Ycf1. Since a dephosphorylated state is unattainable *in vivo*, we mimicked the dephosphorylated state by using the single-site mutants from our ATPase measurements. Consequently, the S908A and S903A mutants showed the most significant deficiency in survival, consistent with their loss of ATPase activity (Figure 4B). The phosphorylation site mutants S869A, S876A, T911A, and S914A all showed no (or slightly elevated) survival in a manner again consistent with ATPase results, leading us to conclude that R-domain driven changes in the ATPase rate are responsible for changes in cadmium detoxification.

### R-domain architecture in the S908A variant most closely mimics dephosphorylated Ycf1

To interrogate changes in the R-domain location and structure in dephosphorylated Ycf1 during glutathione stimulation, we employed a limited proteolysis protection assay (Figure 4C).

Previously, we observed that dephosphorylated Ycf1 leads to a loss of proteolytic fragments corresponding to the R-domain. Of the mutants with nearly complete activity loss, the S908A mutant displayed a digestion pattern most similar to the dephosphorylated form of Ycf1 presented in our previously published study^6^. Missing bands in the digest are within the molecular weight range of a fragment formed by residues 850-930. It is unclear why this banding loss occurred only in S908A and dephosphorylated Ycf1. We initially attributed this effect to increased susceptibility to protease. In light of our cryo-EM structure, we interpret this as a protective effect where the dephosphorylated R-domain resides in the substrate cavity and may be shielded from protease. We further conclude from the similarities in the digest patterns of dephosphorylated and S908A Ycf1 that the observed changes in ATPase rate and cell survival are correlated with changes in R-domain structure/dynamics.

## Discussion

Our dephosphorylated Ycf1 structure reveals a new inactive state that offers insights into the phosphoregulation of ABCC transporters by autoinhibition. The dephosphorylated R-domain packing inside the Ycf1 substrate cavity locks the transporter in the IFwide confirmation. In this state, phosphorylation of the R-domain hairpin bundle, specifically at the S908 position, is likely the master switch that controls the transition of the R-domain from an autoinhibition to activator conformation. In this context, a few mechanisms of autoinhibition in Ycf1 are likely: 1) the R-domain occludes substrate binding directly to prevent transport, 2) the R-domain physically blocks the assembly of the TMDs and, consequently, NBD dimerization forming the power stroke of the ABC transporter conformation switch, 3) the R-domain blocks engagement of the GRD motif^23^, a key motif required for allosteric coupling between NBDs and TMDs, and lastly 4) R-domain sequestration away from the allosteric activator site on NBD1, which our previous phosphorylated Ycf1 structure^6^ showed requires R-domain docking for Ycf1 stimulation. Scenarios 1-3 are antagonistic, whereas scenario 4 represents the loss of a stimulatory effect but not function.

These findings, therefore, suggest a plausible R-domain activation and deactivation cycle (Figure 5). In the resting state, the R-domain maintains the transporter in an inactive conformation. Other groups have predicted this widespread autoinhibitory mechanism to arise from the loss of NBD dimerization by direct binding of the R-domain. In contrast, others suggest a role in the blockage of the substrate pore^9^. Our structure shows that the R-domain impacts both functions. Although phosphorylation at S903 first appears to be the required step in releasing the R-domain from its inhibitory site, we still do not know the order of events leading to subsequent phosphorylation at other R-domain phosphosites. Ultimately, the phosphorylated R-domain hairpin is released from its inhibitory site and partially disassembles its tertiary structure while maintaining its secondary structure to engage the potentiating site on NBD1 near the junction of NBD1, ICL3, and the lasso domain, where the phosphorylated residues S903, S908, T911, and S914 in the R-domain can interact with a cluster of positively charged residues (K615, K617, K655, R716, R770, R206)^6^.

**Figure 5.**
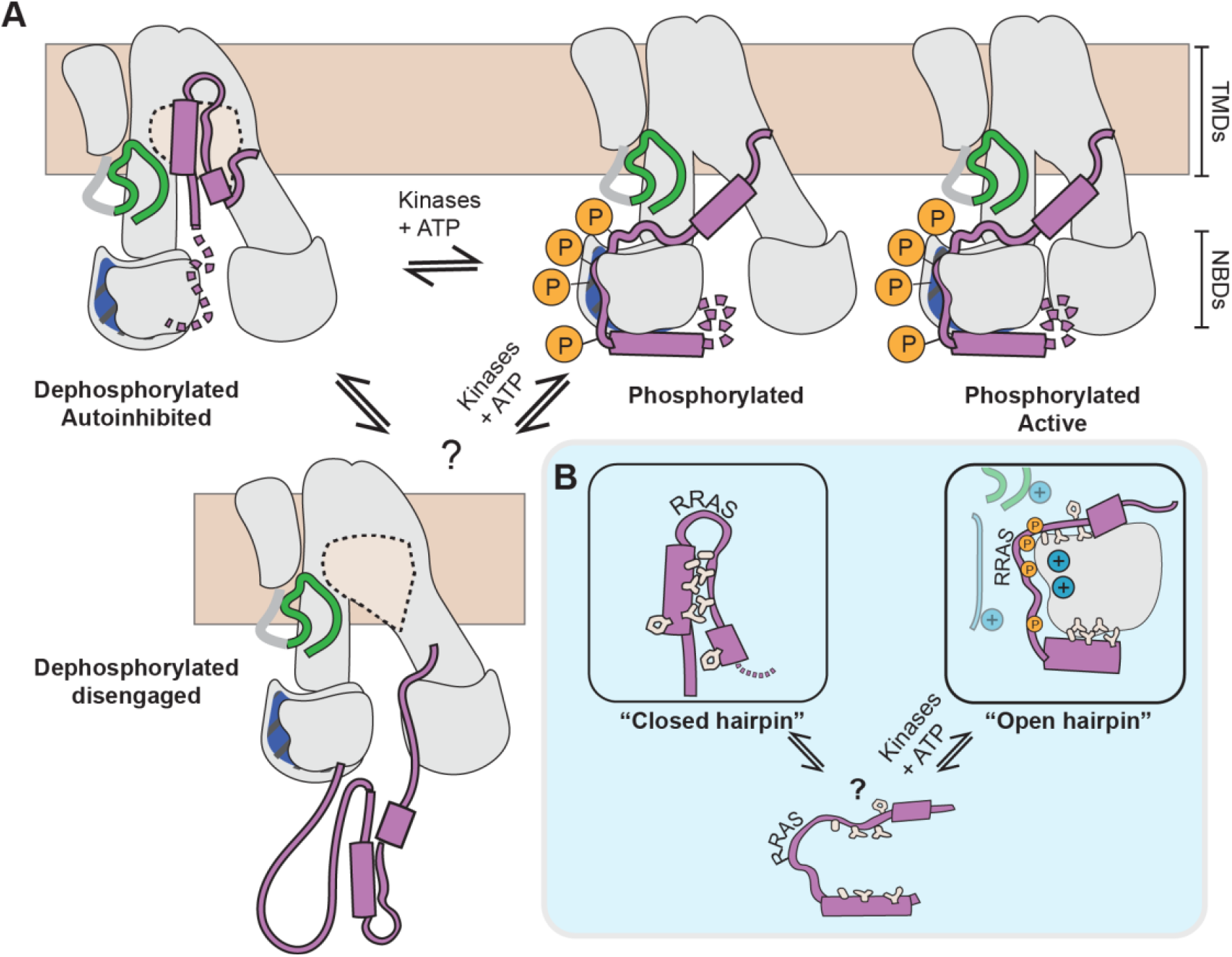
Proposed model for R-domain autoinhibition and activation. **A**. Overall cycle of Ycf1 transitioning from a dephosphorylated state to a phosphorylated state. **B**. The R-domain hairpin loop (purple) is shown with interacting hydrophobic residues (tan). Blue circles represent positive charges near NBD1.

This mechanism is consistent with several previous biochemical results from full-length protein investigations but conflicts with biophysical investigations on isolated R-domain fragments. Solvent accessibility experiments on full-length CFTR using tryptophan fluorescence quenching showed a significant (∼20% or ∼450 residues) change in the solvent-accessible surface area upon phosphorylation consistent with the release of a large domain^22^. In another solvent accessibility experiment using deuterium exchange by attenuated total reflection Fourier transform infrared (ATR-FTIR) spectroscopy, these changes localized to the TMDs^22^. Importantly, these changes are not accompanied by any significant changes in secondary structure. Together, these findings support our hairpin model of transporter autoinhibition in which the unphosphorylated R-domain buries in the substrate cavity, then unwinds upon phosphorylation to release itself from the substrate pocket binding site, and lastly assumes a new position on NBD1 while maintaining its secondary structure. Finally, these experiments show the opposite effect on isolated NBD1^5^, suggesting that the R-domain overall moves from the cavity space to NBD1 and is consistent with our model.

Our data indicate that several R-domain phosphorylation sites only partially affect the ATPase activity and protease accessibility in Ycf1, except for S908A, which mimics the dephosphorylated Ycf1 state and has the most substantial impact on these functions. Patch clamp measurements support this result in an R-domain fragment of CFTR containing S813^25^ (equivalent to S908 in Ycf1) and S798 shown to be required for release of autoinhibition by the R-domain C-terminus in CFTR. Together this suggests that R-domain phosphorylation is the primary driver of activity and the cascade of events needed for transporter activation. However, these findings do not explain kinase recruitment and phosphosite specificity. While Phosphorylated R-domain peptides can partially rescue activity in unphosphorylated CFTR, the loss of its stimulatory effect is seen only in deletion constructs lacking the entire R-domain deletion. Our model reconciles these views and suggests that both allosteric stimulation of NBD1 and relief of autoinhibition by the R-domain are parts of the same R-domain activation mechanism where release from the substrate cavity precedes/primes an NBD reorganization to stimulate activity.

The exact sequence of events exposing S908 to kinases remains a mystery in the context of our structure because S908 is not accessible to kinases in the bound state we observe. S908 accessibility could be a stochastic process or could be driven by the phosphorylation of a secondary site, helping explain their partial phenotype. One possibility is that the R-domain position seen in CFTR structures is another state showing these “released” states are accessible for phosphorylation. It is likely that once a critical mass of phosphorylation is achieved in such a small volume, electrostatic repulsion disfavors hairpin formation and instead favors engagement with a positively charged surface of NBD1, consistent with phosphomimetic mutation experiments suggesting negative charge as sufficient cause for R-domain rearrangement^24^.

Multiple factors suggest that an activation cycle dependent on phosphorylation/dephosphorylation may be conserved across several ABCC family members. The R-domain C-terminus maintains a higher degree of conservation between the several ABCC members, including CFTR, MRP1, and Ycf1. In members with multiple phosphorylation sites, the relative spacing between phosphosites is also conserved, especially in proximity to the S908 site, which has been considered important even when sequence conservation is not high (Figure S3). Though these transporters maintain different functions, MRP1 and Ycf1 can functionally replace one another, and all three have been reported to have glutathione transport activity^4,14^. Finally, recent investigations using single-molecule fluorescence resonance energy transfer (smFRET) on CFTR suggest phosphorylation governs NBD dimerization for chloride conductance^26^.

The substrate cavity interactions in the autoinhibited R-domain conformation are also well-conserved among many ABCC members. The dephosphorylated R-domain makes several contacts synonymous with glutathione binding in MRP1^8^. The R-domain architecture shares a striking resemblance to the IC47 viral peptide inhibitor from herpes simplex virus bound to the TAP1/TAP2 transporter (Figure S4), which adopts a similar hairpin architecture stabilized by a hydrophobic zipper and electronegative residues within the hairpin loop^27^. Other ABC transporters families with similar linkers and controlled by phosphorylation may have similar mechanisms. ABCB1 (P-glycoprotein) is suggested to have a helical hairpin structure in its NBD1-TMD2 linker based on similarity to excreted WXG motif hairpin proteins (Figure S4) that are secretion substrates for Type VII type ABC secretion system^9^. In P-glycoprotein, this proposed model similarly places phosphorylated serines at the apex of the hairpin, similar to the dephosphorylated Ycf1 structure.

In conclusion, our dephosphorylated Ycf1 structure shows a dramatic rearrangement of the R-domain that occludes the transporter substrate binding pocket, resembling the peptide-bound state of TAP1/TAP2. It suggests a mechanism where the R-domain blocks activity by binding substrate, by preventing NBD dimerization, and by engaging with allosteric elements that communicate ATP binding to the substrate cavity. Lastly, our data support S908 as a dominant driver of Ycf1 activation, likely through the release of the R-domain hairpin inhibition and by direct stimulation of NBD1.

## Method

### Ycf1 expression and purification

The *S. cerevisiae* Ycf1 expression construct used here is the same as in our previous study^6^. R-domain phosphorylation site mutants were constructed by site-directed mutagenesis and verified by sequencing (Elim Biopharmaceuticals, Inc). Protein was expressed and purified as previously described^6^. Briefly, the DSY-5^38^ strain of *S. cerevisiae* (Genotype MATa leu2 trp1 ura3-52 his3 pep4 prb1) was transformed with wild-type or mutant Ycf1-encoding plasmids and selected on YNB-His agar medium plates. Transformants were inoculated into a primary culture of 50 mLYNB-His media (yeast nitrogen base with ammonium sulfate 0.67% w/v, glucose 2% w/v, CSM-His 0.077% w/v) and grown for 24 hours at 30°C with 200 rpm shaking. Then, 15 mL of the primary culture was used to inoculate 750 mL of a secondary YNB-His culture and grown similarly to the primary culture for an additional 24 hours. Protein expression was induced by adding 250 mL of 4X YPG media (4% w/v yeast extract, 16% w/v peptone, and 8% w/v galactose). The culture was grown for an additional 16 hours. Finally, cells were collected by centrifugation at 5000xg and 4°C for 30 minutes and stored at -80°C.

Crude membranes were prepared by cells resuspended in lysis buffer (50 mM Tris-Cl pH 8.0, 300 mM NaCl, protease inhibitor cocktail mix) and subject to bead beating as previously described^6^. Briefly, the lysate was separated from beads by filtering through a coffee filter, and membranes were harvested by ultracentrifugation of the clarified lysate at 112,967xg for 1.5 hours before storage at -80°C. Membranes were solubilized for 4 hours at 4°C at a ratio of 15 mL resuspension buffer (50 mM Tris-Cl pH 7.0, 300 mM NaCl, 0.5% 2,2-didecylpropane-1,3-bis-β-D-maltopyranoside (LMNG)/0.05% cholesteryl hemisuccinate (CHS) supplemented with protease inhibitor cocktail) per gram of membrane. The insoluble fraction was separated by centrifugation at 34,155xg for 30 min at 4°C followed by filtration of the supernatant through a 0.4 µM filter prior to loading onto a 5 mL Ni-NTA immobilized metal affinity chromatography (IMAC) column (Bio-Rad) equilibrated in buffer A (50mM Tris-Cl, 300mM NaCl, 0.01% LMNG/0.001% CHS, pH 7.0).

IMAC immobilized protein was purified by first washing with 50mL (10 column volume (CV)) of buffer A followed by a gradient of buffer B (50 mM Tris-Cl, 300 mM NaCl, 500 mM Imidazole 0.01% LMNG/0.001% CHS, pH 7.0) using the following steps sizes: 10 CV of 6% Buffer B, 2 CV of 10% Buffer B, 2 CV of 16% Buffer B and 2 CV mL of 24% Buffer B. Protein was eluted with 4-5CV of 60% buffer B and diluted immediately with 10-fold buffer A prior to being concentrated at 3095xg at 4°C in a 100 kDa cutoff Amicon concentrator (Millipore). The process was repeated three times to remove the excess imidazole prior to sample injection onto a size exclusion Superose 6 Increase 10/300 GL column (GE Healthcare) equilibrated in SEC buffer (50 mM Tris, 300 mM NaCl, pH 7.0) supplemented with 0.01% LMNG/0.001% CHS (Figure S5). Protein concentration was measured by Bicinchoninic acid assay (Pierce) and used immediately, without freezing, for biochemical and structural analyses.

Ycf1 dephosphorylation was performed using our previously established method^6^. SEC-purified sample Ycf1/Ycf1-E1435Q (5 µg) was treated with 1 µL of Lambda phosphatase (Lambda PP, NEB) for 1hr at 30ºC. Following treatment, the sample was subjected to a second round of SEC purification, the same as above, to remove the phosphatase. For cryo-EM samples, SEC purification was performed in SEC buffer containing 0.06% digitonin, concentrated, quantified, and immediately used for grid freezing.

### ATPase activity assay

The ATP hydrolysis activity of purified Ycf1 was measured at 30ºC using the enzyme-coupled assay as described previously^6^. Reaction samples were 75 µL each and contained 6.45 µg of purified protein in a reaction mixture consisting of 20 mM Tris-HCl pH 7.0, 10 mM MgCl_2_, 1 mM PEP, 55.7 / 78.0 U/mL PK/LDH, 0.3 mg/mL NADH and 1 mM ATP. Substrate-induced ATPase activity was measured by adding 8 or 16 µM GSSG to the reaction mix. ATPase rates were calculated as a function of NADH consumption measured as the change in absorbance at 340 nM for 45 min on a Synergy Neo2 Multi-mode Microplate Reader (BioTek). ATPase rates were calculated using linear regression in GraphPad Prism 9.

### Yeast survival spot assay

A Ycf1-KO *S. cerevis*iae strain transformed with wild-type or mutant Ycf1 was grown overnight on YNB-His agar plates. The following day, a dilution of the overnight culture with an OD600 reading of 0.1 was prepared in 0.9% saline. Five-fold serial dilutions were prepared and 5 μL of each spotted onto YRG (yeast nitrogen base with 0.67% w/v ammonium sulfate, 1 % w/v raffinose, 2% w/v galactose, 0.077% w/v CSM-His, and 2% w/v agar) agar plates with and without 150 µM CdCl_2_. Plates were incubated for five days at 30ºC and photographed using a Bio-Rad Chemidoc MP Imaging System. The assay was performed in quadruplets (two biological replicates, with two technical replicates per biological replicate). Representative images are shown in Figure 4.

### Limited trypsin proteolysis

Ycf1 samples were treated with trypsin from bovine pancreas (Sigma) at different concentrations (0, 4, 8, and 20 µg/mL) for 1h on ice in a 20 µL reaction volume containing 6 µg of purified wild-type or mutant Ycf1. After 1h, the reaction was stopped by adding 1 μL of soybean trypsin inhibitor (Sigma) from a stock at 1 µg/mL concentration and incubating for an additional 15 min on ice. Gel loading samples were prepared by mixing 5 µL of each reaction with 1X SDS loading dye containing 100 mM DTT and separated on 4–20% SDS-PAGE gels (Bio-Rad) before staining with Coomassie Brilliant Blue R-250 stain. Experiments were performed at least in triplicate.

### Dephosphorylated Ycf1 grid preparation and cryo-EM data collection

5 µL of dephosphorylated Ycf1-E1435Q at 10.56 mg/mL were applied on QF-1.2/1.3-4Au grid (Electron Microscopy Sciences) using a Leica EM GP2 set to 10ºC and 80% humidity. Following sample application, the grid was incubated for 10s followed by a 3.5 second blot on Whatman 1 paper and immediately plunge-frozen in liquid ethane equilibrated to -185ºC.

A total of 5,904 movies were recorded on a Titan Krios at 300 kV equipped with a K3 Summit detector (Gatan) using SerialEM v4.0.0beta software at the Pacific Northwest Center for Cryo-EM (PNCC). Data were collected in super-resolution mode at 81,000X magnification with a defocus range of -0.6 to -1.9 µm. Each movie contains 75 frames with a total electron dose of ∼52 electrons /Å^2^.

### Cryo-EM data processing

The dephosphorylated Ycf1-E1435Q dataset was processed starting with Relion 4.0. Image stacks with a pixel size of 1.069 Å/pixel generated from drift correction using MotionCor2^30^ in Relion 4.0 followed by contrast transfer function (CTF) estimation using CTFFIND4.1^31^. Micrographs with estimated CTF resolutions of 4.5 Å or better were selected (5,152 micrographs) and subject to template-free manual particle picking and 2D classification. These 2D classes were then used as templates for automated particle picking, and a total of 5,424,196 particles were picked and extracted with 4X binning to 4.277 Å/pixel and a box size of 240 pixels (Figure S1A). Three rounds of 2D classification were performed, resulting in 1,341,751 particles subjected to ab-initio 3D map generation. Using the ab-initio 3D map as a reference, two rounds of 3D classification were performed. From the second round of 3D classification, the dominant class (class 3, 46.7% particles) was selected and particles were unbinned to a pixel size of 1.069 Å/pixel and extracted with a 240 pixel size box. The unbinned particles were further subject to 3D refinement, followed by 3D classification without alignment. The best class out of three was selected based on resolution, and particles were re-extracted to a box size of 360 pixels at a size 1.069 Å/pixels. 3D refinement was then performed in RELION using SIDESPLITTER^3^. The refined class was subjected to iterative rounds of CTF refinement, Bayesian polishing, and postprocessing in RELION. A final round of alignment-free 3D classification was performed to remove structural heterogeneity followed by one round of Bayesian polishing, 3D refinement, and postprocessing in RELION, resulting in a 3.1Å resolution map.

### Ycf1 model building

A Ycf1 model from AlphaFold2^37^ was used as an initial model combined with elements of our previous Ycf1 structure^6^. The Isolde^36^ plugin for ChimeraX^34^ and manual fitting in COOT^32^ were used for model improvement. To interpret regions of the R-domain with weak density, two alternate approaches to generate locally sharpened maps, Phenix Density Modification^39^ and DeepEMhancer^40^, were compared but were not used for model refinement in Phenix^33^ (Supplementary feigure 2). Only regions of the model that were visible in the original Relion maps were kept and iterative cycles of real-space refinement were performed by Phenix^33^ against only the original Relion Postprocess map. B-factor blurred (ie positive B-factor applied) Relion maps were also generated in COOT and used to trace the R-domain. Alternate traces of the R-domain in opposite directions were extensively tested using the per residue Correlation Coefficient in Phenix refinement as a guide. All statistics are reported against the Relion Postprocess map (Table 1). Figures were prepared using UCSF ChimeraX^34^. R-domain and ligand binding analysis was performed using the software Ligplot^41^.

**Table1.**
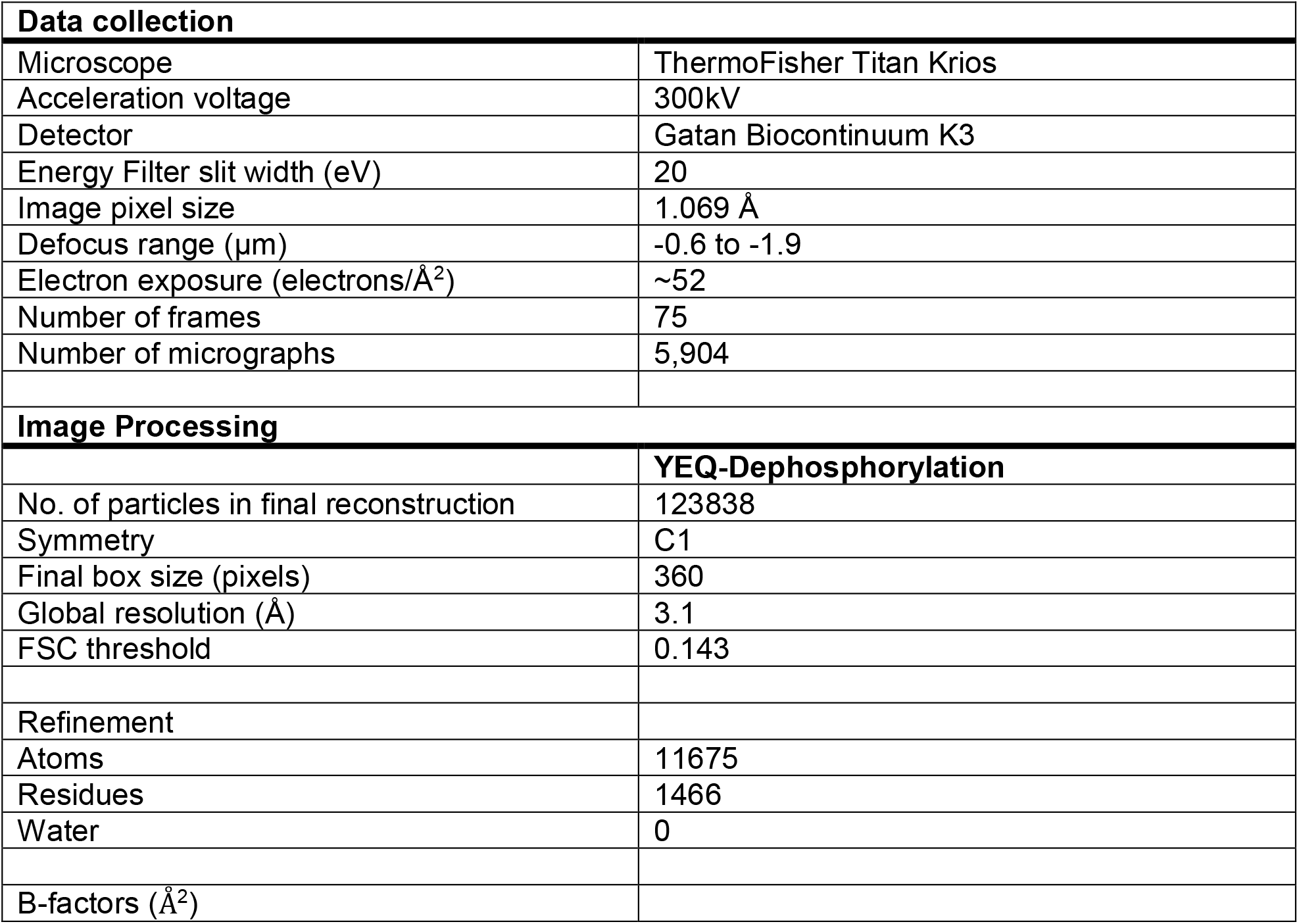
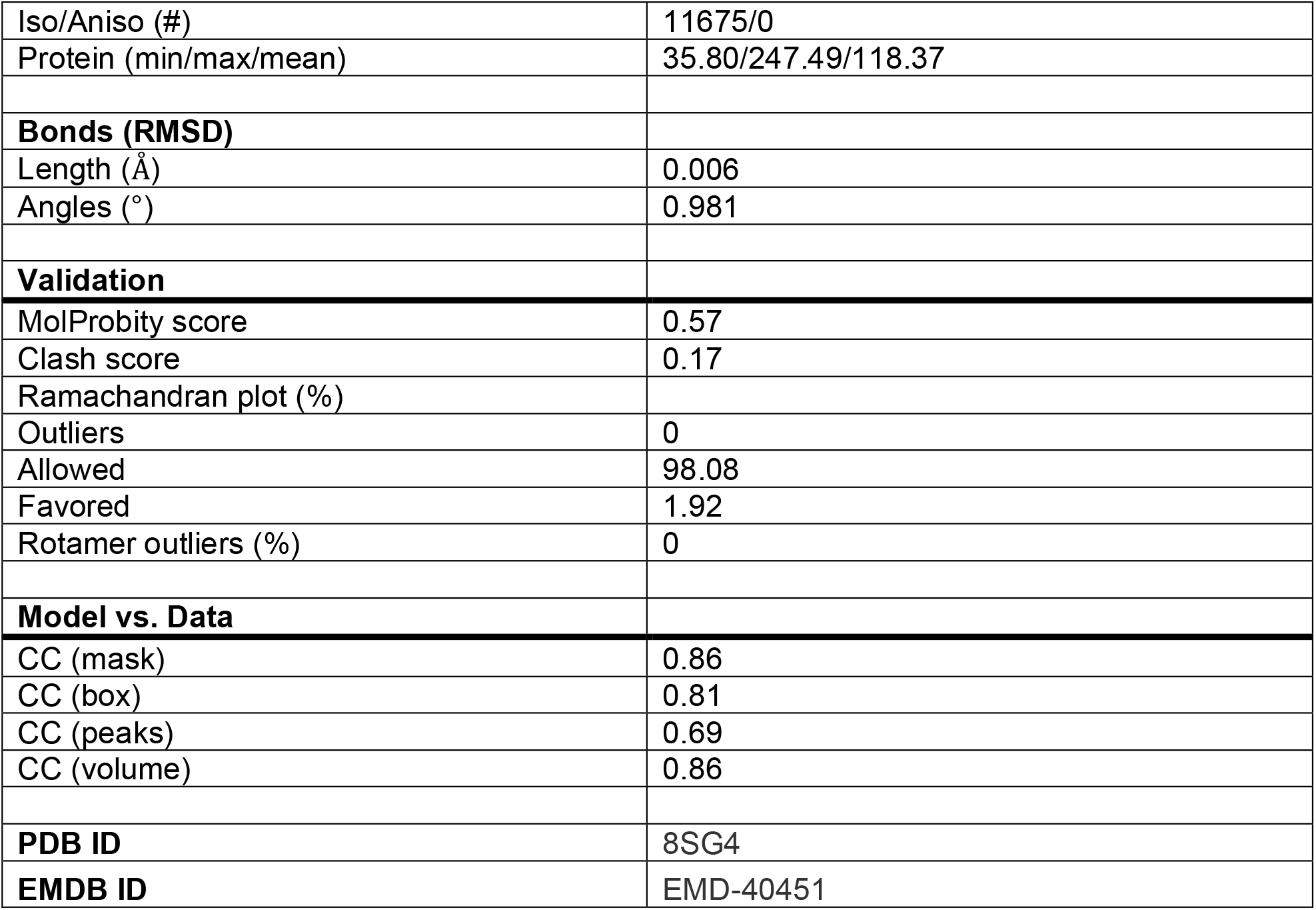
Cryo-EM data collection and refinement statistics.

## Author contributions

Conceptualization: TMT; Methodology: NKK, TMT; Investigation: NKK; Visualization: NKK, TMT; Funding acquisition: TMT; Supervision: TMT; Writing – original draft: NKK, TMT; Writing – review & editing: NKK, TMT.

## Acknowledgments

A portion of this research was supported by NIH grant U24GM129547 and performed at PNCC at OHSU and accessed through EMSL (grid.436923.9), a DOE Office of Science User Facility sponsored by the Office of Biological and Environmental Research. This work was also supported by grants from the National Institute of Allergy and Infectious Disease (NIH R01 AI156270 (TMT)), NIH S10 OD011981 (Life Sciences North Imaging Facility at the University of Arizona) and the University of Arizona BIO5 Postdoctoral Fellowship Award (NKK). We also thank the staff at the Pacific Northwest Center for Cryo-EM (PNCC), especially Nancy Meyer, for assistance with data collection. We also thank Tarjani M. Thaker for critical reading of the manuscript, Meghna Gupta for assistance with data interpretation, and members of the Tomasiak lab for helpful discussions.

