## Supplementary Figures for "Structural Basis for Autoinhibition by the Dephosphorylated Regulatory Domain of Ycf1": Biorxiv-sup.pdf

**Supplementary Figure 1**

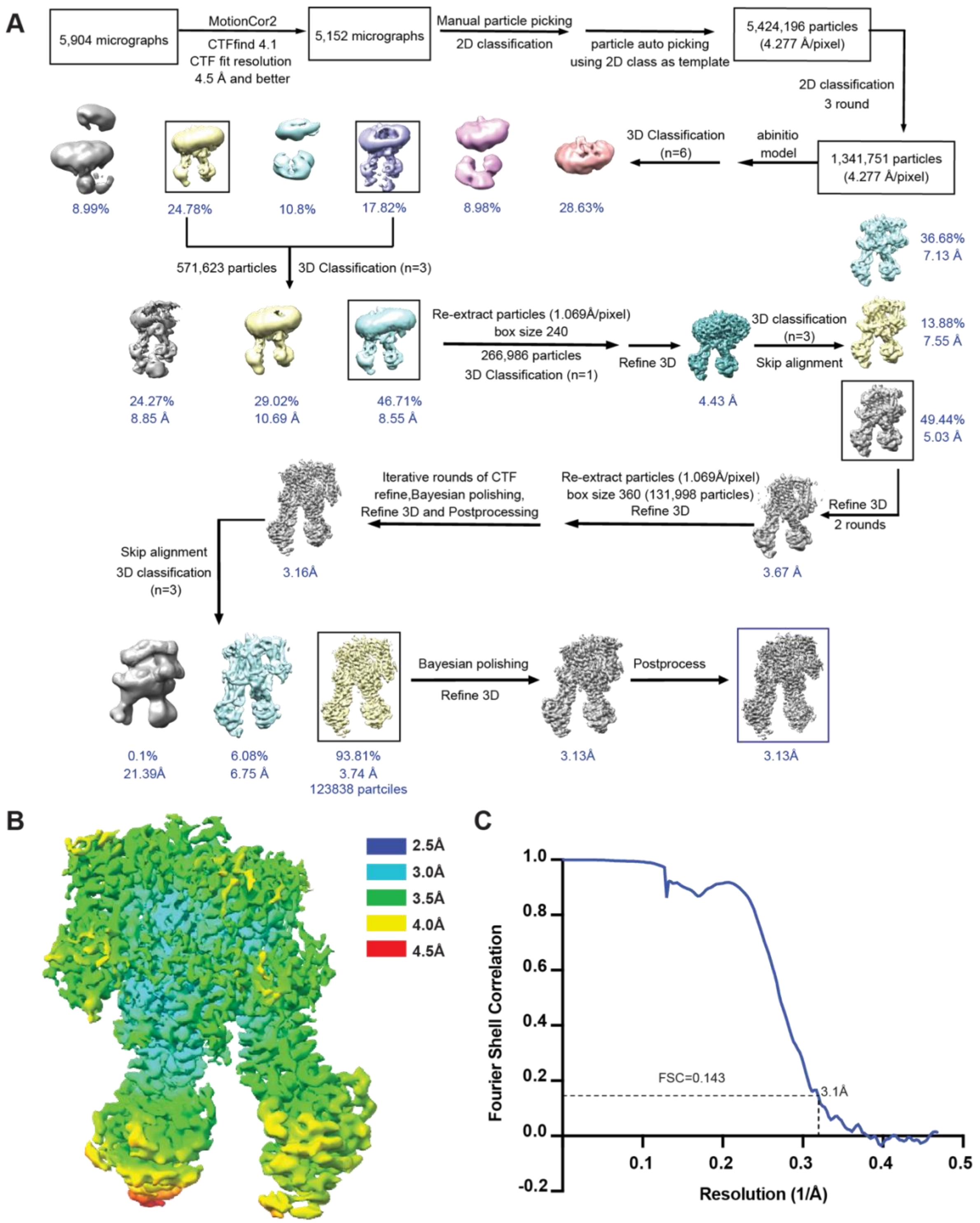


**Figure S1: Cryo-EM data processing workflow of dephosphorylated Ycf1. A**. Data processing workflow for dephosphorylated *S. Cerevisiae* Ycf1-E1435Q. **B.** Local resolution estimation of the final map. **C.** Fourier shell correlation curve with resolution cutoff of 0.143.

**Supplementary Figure 2**

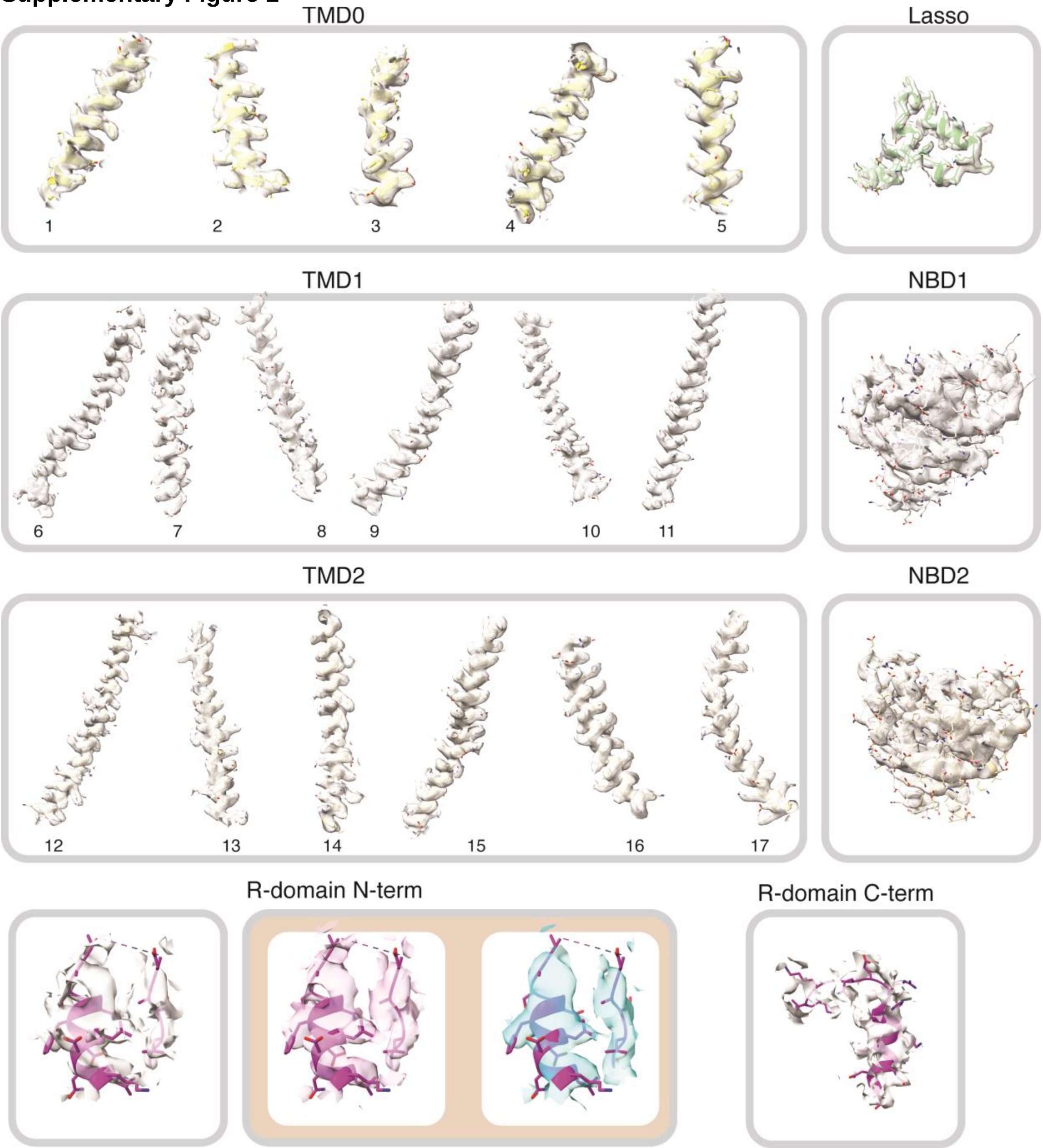


**Figure S2: Cryo-EM densities for secondary structure elements and domains of dephosphorylated Ycf1.** Cryo-EM density corresponding to secondary structural elements and domains observed in the dephosphorylated Ycf1- E1435Q structure. R-domain density is shown for the N- and C-termini. The tan box shows alternate maps of the R-domain n-terminus used to help trace the density including left phenix density modification[39] (left, pink) and DeepEMhancer[40] (right, blue). The cryo-EM density in the TMDs, NBDs, Lasso domain are contoured to a threshold level of ~0.01 and the R-domains N-termini are contoured to ~0.007 and the R-domain C-termini to ~0.006. The color scheme is identical to main text Figure 1.

**Supplementary Figure 3**

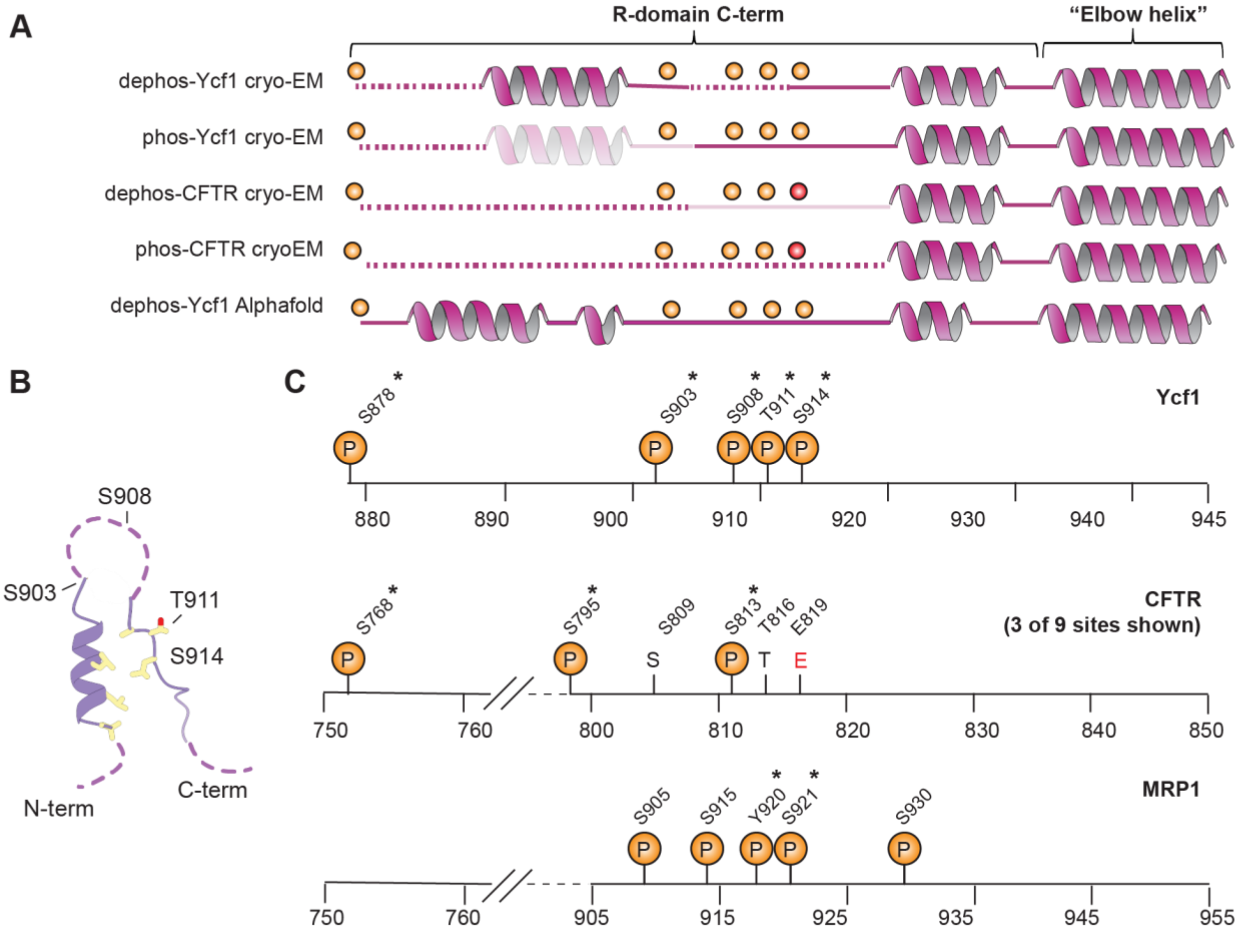


**Figure S3: Comparative analysis of secondary structural features of the R-domain. A.** Topology of secondary structure in the R-domain C-terminus of Ycf1 or CFTR determined from cryo-EM structures[6,7] or predicted by AlphaFold2[42], with corresponding phosphorylation sites highlighted with an orange circle. Red circles denote the presence of a glutamate residue where a phosphorylation site exists in the other proteins. Secondary structure shown represents the structurally conserved regions observed in the respective cryo-EM maps, whereas the transparent helix in Ycf1-phos denotes a region for which cryo-EM density was weak and a model could not be built. **B.** Closeup of the Ycf1 R-domain hairpin loop architecture and relative arrangement of phosphorylation sites. Hydrophobic residues forming key interactions are shown as sticks. **C.** Position of phosphorylation sites, conserved residues, and negatively charged amino acids in the R-domain of Ycf1 (top), CFTR (middle), and MRP1 (bottom). Asterix (*) denotes sites to be confirmed to be phosphorylated.

**Supplementary Figure 4**

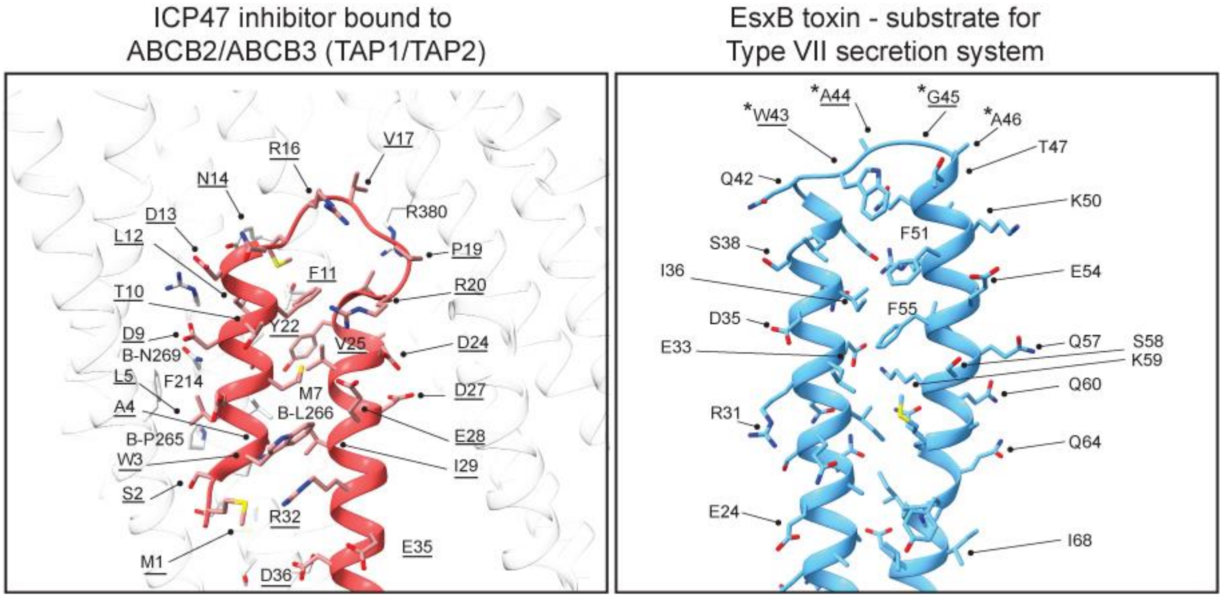


**Figure S4: R-domain-like architecture of peptide ligands in an ABC transporter or transporter with ABC-like ATPase domain. A.** Specific interactions between the herpes simplex virus ICP47 peptide inhibitor (red cartoon) in a closed hairpin loop formation bound to TAP1/TAP2 (grey cartoon; PDB ID:5U1D[27]). Underlined residues denote those on the ICP47 peptide itself. **B.** Alphafold2 structure of the type VII secretion system substrate peptide EsxB from *Bacillus antracis* in a hairpin loop form. The symbol (*) denotes residues of the WxG motif.

**Supplementary Figure 5**

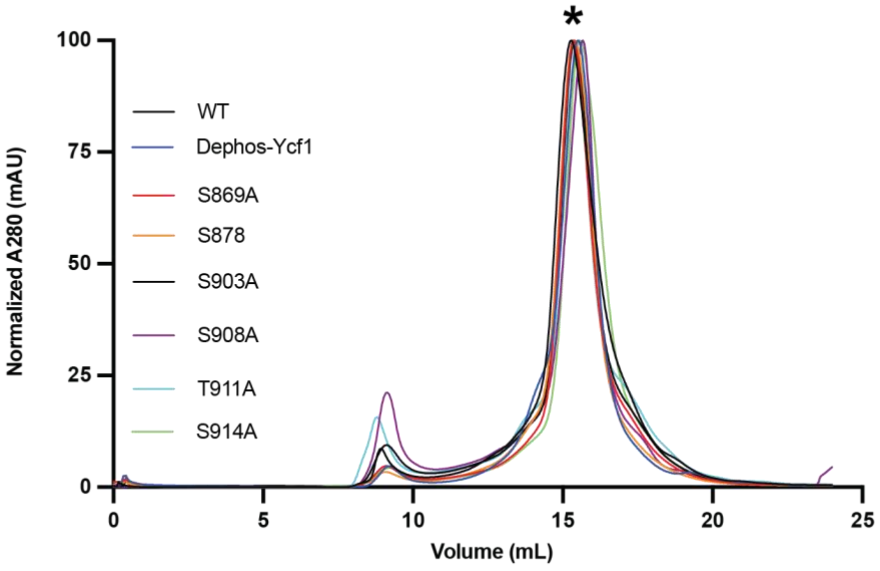


**Figure S5: Size exclusion purification profile of Ycf1 and mutant variants.** Chromatograms are representative of mutants of Ycf1 purified using a Superose 6 Increase 10/300 GL Column (Cytiva).
